# Pathogenic tau disrupts the cellular program that maintains neuronal identity

**DOI:** 10.1101/2021.03.05.434166

**Authors:** Adrian Beckmann, Paulino Ramirez, Maria Gamez, William J. Ray, Bess Frost

## Abstract

Neurons in human Alzheimer’s disease acquire phenotypes that are also present in various cancers, including over-stabilization of the cytoskeleton, nuclear pleomorphism, decondensation of constitutive heterochromatin, and aberrant activation of the cell cycle. Unlike in cancer, in which cell cycle activation drives tumor formation, activation of the cell cycle in post-mitotic neurons is sufficient to induce neuronal death. Multiple lines of evidence suggest that abortive cell cycle activation is a consequence of pathogenic forms of tau, a protein that drives neurodegeneration in Alzheimer’s disease and related “tauopathies.” We have combined network analysis of human Alzheimer’s disease and mouse tauopathy with mechanistic studies in *Drosophila* to discover that pathogenic forms of tau drive abortive cell cycle activation by disrupting the cellular program that maintains neuronal identity. Mechanistically, we identify Moesin, a prognostic biomarker for cancer and mediator of the epithelial-mesenchymal transition (EMT), as a major effector of tau-induced neurotoxicity. We find that aberrant activation of Moesin in neurons acts through the actin cytoskeleton to dysregulate the cellular program that maintains neuronal identity. Our study identifies mechanistic parallels between tauopathy and cancer and sets the stage for novel therapeutic approaches.

## Introduction

Post-mitotic cells such as neurons require persistently active cellular controls to maintain a quiescent, non-cycling, state of terminal differentiation^1–3^. A curious aspect of postmortem human Alzheimer’s disease brains as well as brains of multiple animal models of Alzheimer’s disease and related tauopathies is the neuronal upregulation of proteins that are associated with cell cycle activation^4,5^. Unlike cancer, in which uncontrolled cell division causes tumor formation, cell cycle activation in post-mitotic neurons causes neuronal death rather than neuronal division^6–9^. Cellular phenotypes beyond cell cycle activation are shared between tauopathy and various cancers, including over-stabilization of the actin cytoskeleton^10–13^, changes in nuclear shape and the lamin nucleoskeleton^14,15^ and loss of heterochromatin-mediated transcriptional silencing^16,17^ all of which are also known to be important determinants of cellular identity^18–20^.

Some basic biological functions require dynamic shifts between programs that control cellular identity and those that promote cellular plasticity. During EMT, for example, transdifferentiation of epithelial cells into mesenchymal cells is important for wound healing^21,22^ and organ development^23^. Mechanistically, the cytoskeletal remodeling that occurs with EMT causes breakdown of cell-to-cell connections and depletion of proteins that maintain a terminally differentiated epithelial identity. A shift from cellular identity to cellular plasticity can also mediate disease. In cancer, for example, EMT disrupts the terminally differentiated epithelial phenotype to facilitate tumor metastasis^24,25^, cell cycle activation and consequent malignancy^26,27,28^.

In mature neurons, a terminally differentiated state is maintained by “terminal neuronal selector proteins,” key transcription factors that are in part regulated by the extracellular environment^29–32^. Based on the parallels between cellular phenotypes in cancer and tauopathy, as well as the relationship between cancer and disrupted cellular identity, we hypothesized that pathogenic forms of tau interfere with the cellular program that maintains neuronal identity.

In this study, we first analyzed transcriptional networks in human and mouse tauopathy to gain insights into the relationships between co-expressed genes and their association with biological determinants of cellular identity. These analyses identified *Moesin*, which is well-known for its role in EMT and cancer metastasis^13,26^, as a highly connected gene in a co-expression module that is related to EMT and cancer in both human Alzheimer’s disease and across disease stages in tau transgenic mice. We next leveraged the genetic power of *Drosophila* to discover that features of EMT are present in brains tau transgenic *Drosophila*, that aberrant Moesin activation mediates tau-induced overstabilization of the actin cytoskeleton and consequent abortive cell cycle activation, and that activation of Moesin in post-mitotic neurons disrupts the cellular program that maintains terminal neuronal differentiation. Overall, our study suggests that pathogenic tau causes abortive cell cycle activation in neurons by deleteriously mimicking the cellular processes driving EMT.

## Results

### Network analysis of postmortem human Alzheimer’s disease brains identifies a large module related to EMT and cancer

Limitations of RNA-seq-based differential gene expression analysis are the inability to understand the relationships between expressed genes and to stratify genes in a biologically meaningful manner. Weighted Gene Correlation Network Analysis (WGCNA) uses transcriptomic data to analyze relationships between co-expressed genes, cluster groups of highly co-expressed genes into modules, and identify the “hub genes” within each co-expression module^33^. To begin to test the hypothesis that pathogenic tau disrupts a biological program that maintains cellular identity, we performed WGCNA using publicly available RNA-seq data from temporal cortex of postmortem human control (n=57) and late-stage Alzheimer’s disease (n=82) patients generated by the Accelerating Medicines Partnership – Alzheimer’s Disease (AMP-AD). Co-expression network analysis identified four distinct groups, or modules, of highly co-expressed genes within the human dataset (Fig. 1A-C).

**Figure 1.**
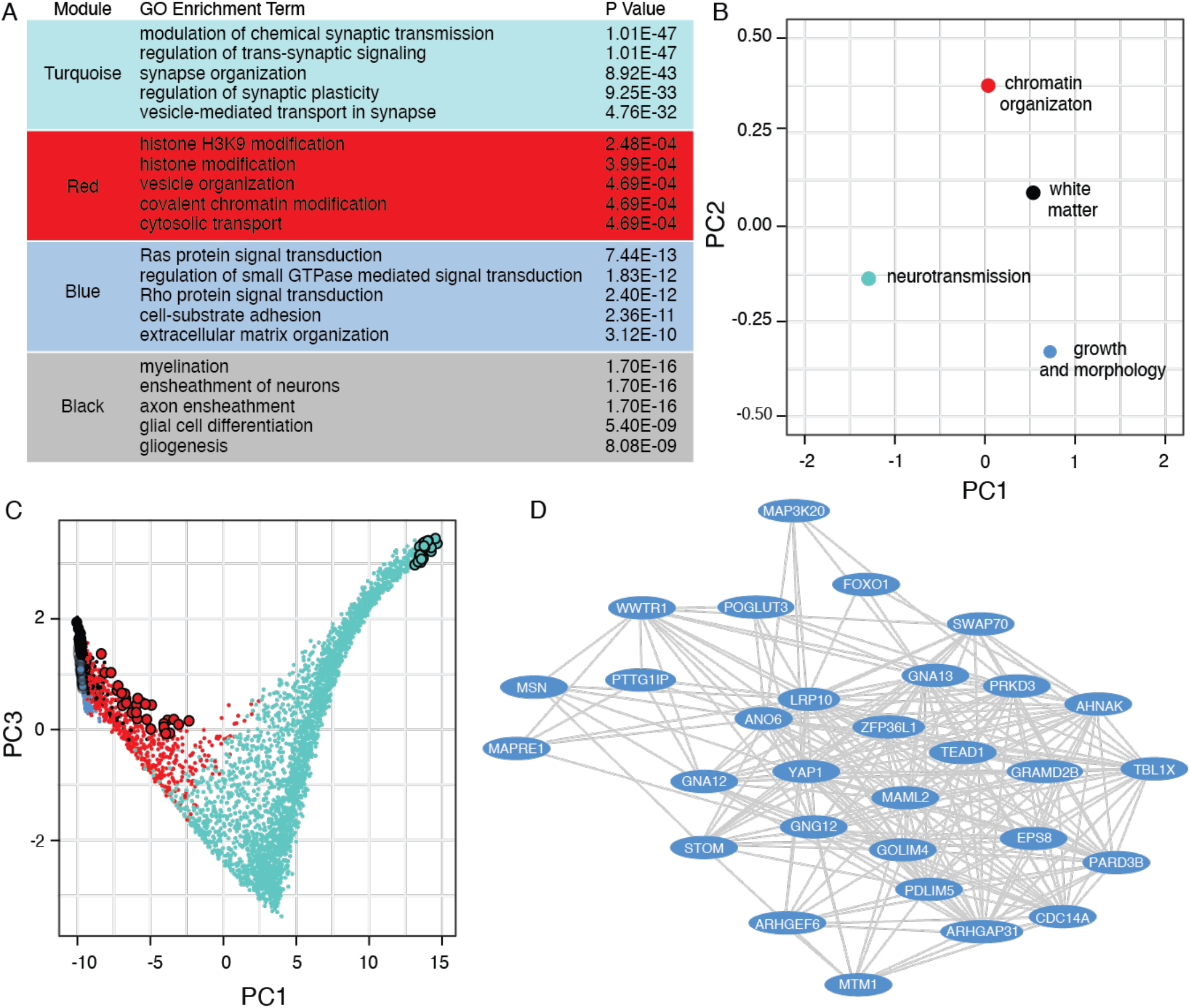
Genes associated with malignancy are systematically upregulated in human Alzheimer’s disease brains, while genes associated with neurotransmission are systematically downregulated. **(A)** Biological processes with the highest degree of significant enrichment based on Gene Ontology. The full table of enriched processes are provided in Supplementary Table 1. **(B)** Multidimensional scaling plot of the first and second principal components for module eigengenes identified by WGCNA. **(C)** Multidimensional scaling plot of the entire human network using principal component three as a function of principal component one. Each point is a single gene. Larger points represent hub genes, which ranked as the top 30 most connected genes for each module. **(D)** Hub genes of the blue module. Each oval represents a node while each line represents the weighted connection between nodes. Colors within each figure correspond to the module assignment for each group of co-expressed genes.

For each module, we performed biological enrichment analysis using Gene Ontology^34,35^ (Fig. 1A, Table S1) and gene-disease association using DisGeNET^36^ (Table S2). In support of our overall hypothesis that pathogenic tau disrupts a biological program that maintains cellular identity, we found that the blue module, which consists of 600 genes, was associated with cellular processes including EMT, organ morphogenesis and cytoskeletal organization (Fig. 1B, 1C, Table S1), and various malignancies such as glioblastoma multiforme and invasive breast carcinoma (Fig. S1A). Many blue module hub genes, such as (*MSN)*, *YAP1*, *TEAD1*, and *WWTR1* are well-known for their role in EMT and cancer^37–40^ (Fig. 1D).

Module eigengenes, defined as the first principal component of a module, can be used to measure the degree of similarity between modules in a network^41^. Our analysis revealed a negative association between the blue module and the turquoise module, which was enriched for biological processes associated with neurotransmission and neurological disorders (Fig. 1B, C, Table S1, S2). Most differentially expressed genes of the blue module were upregulated, while most differentially expressed genes of the turquoise module were downregulated (Fig. S1B, S1C). As neurotransmission is a fundamental function of neurons and an indicator of a mature neuronal identity^32,42^, our human network analyses suggest that brains of patients with late-stage Alzheimer’s disease undergo a systematic downregulation of biological processes associated with neuronal identity (turquoise module) that is accompanied by a systematic upregulation of biological processes associated with EMT and cancer (blue module).

### Age-dependent network analysis of tau transgenic mice identifies biological processes in human Alzheimer’s disease that are driven by pathogenic tau and are conserved across disease stage

Human Alzheimer’s disease is neuropathologically defined by the presence of amyloid β plaques and neurofibrillary tau tangles^43,44^. Limitations of a gene expression network constructed from late-stage postmortem human Alzheimer’s disease brain tissue are 1) the presence of co-pathologies, which do not allow one to differentiate between changes that are a specific consequence of amyloid β, pathological forms of tau, or other events such as vascular damage, and 2) the inability to determine how co-expression networks change as the disease progresses. To determine the specific consequences of pathological tau on gene expression networks and to identify changes that are conserved across disease state, we constructed a co-expression network using RNA-seq data from brains of three, six, and nine-month-old rTg4510 tau transgenic mice. This model features transgenic *CaMKIIa*-driven forebrain expression of the human *MAPT* (tau) gene carrying the disease-associated *P301L* mutation^45,46^ (referred to hereafter as “tau transgenic mice” for simplicity).

Similar to our human network analysis, the mouse network featured a negative correlation between modules associated with neurotransmission/neuronal identity, and modules associated with growth and development (Fig. 2A-D). We also found that the majority of differentially expressed genes in the neurotransmission-related turquoise module were significantly downregulated, while the majority of differentially expressed genes in the blue and yellow modules, which were associated with immune response, growth, and morphology, were significantly upregulated (Fig. S2). Importantly, these analyses also suggested that downregulation of the neurotransmission-related turquoise module was not a simple consequence of neuronal loss, as the network was composed of both control and tau transgenic mice at early, mid, and late-stage disease (Fig. S2).

**Figure 2.**
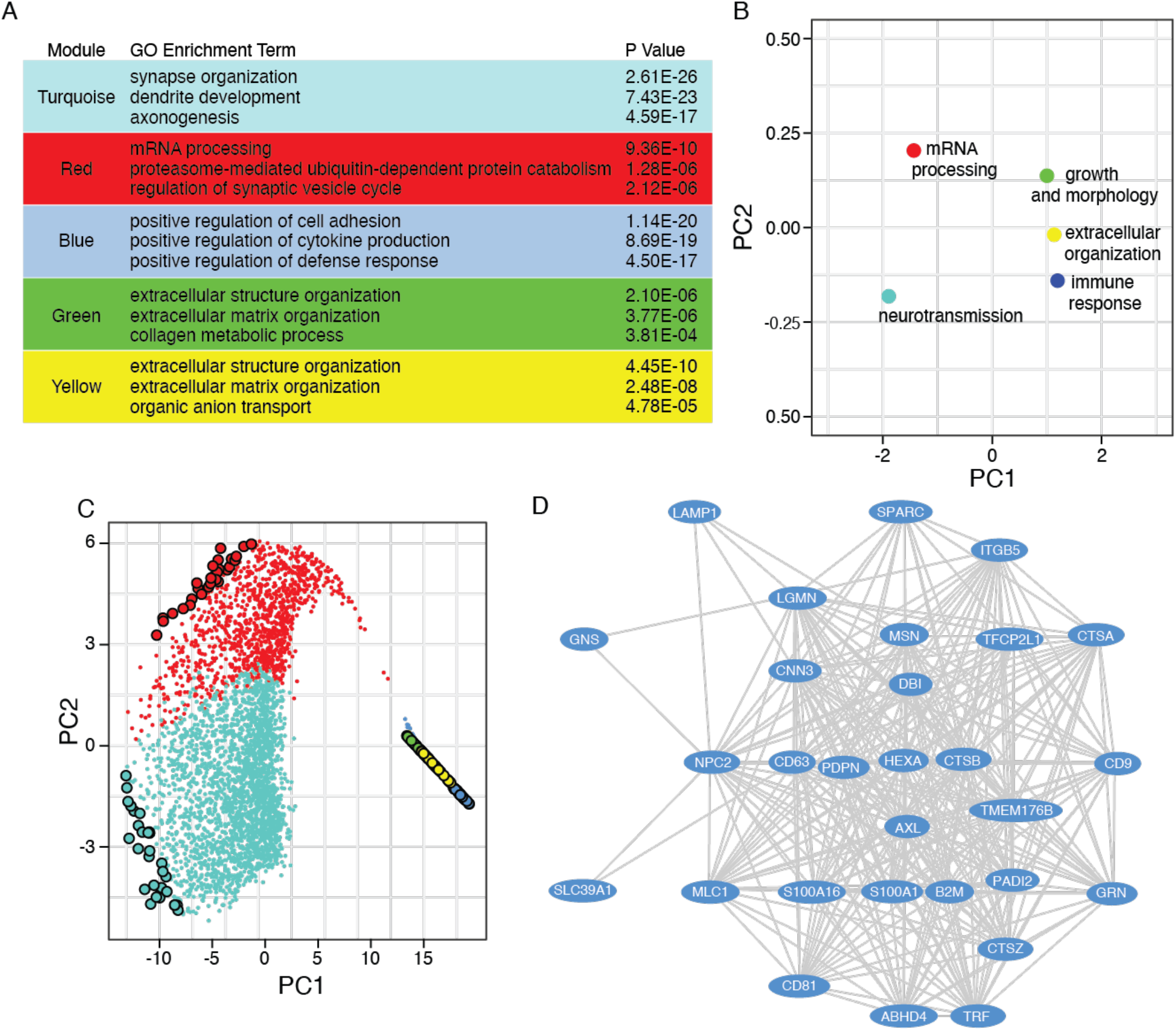
Network analysis tau transgenic mice across aging reveal pathogenic tau drives key pathway signatures present in human Alzheimer’s disease brains. **(A)** Table showing the three most significantly enriched terms for each module based on Gene Ontology. **(B)** Multidimensional scaling plot of the first and second principal components for module eigengenes identified by mouse WGCNA. **(C)** Multidimensional scaling plot of the entire mouse network using principal component three as a function of principal component one. Each point is a single gene. Larger points represent hub genes which ranked as the top 30 most connected genes for each module. **(D)** Hub genes of the blue module. Each oval represents a node while each line represents the weighted connection between each node. Colors within each figure correspond to the module assignment for each group of co-expressed genes.

We next asked whether the modular structure of the human network was preserved in the mouse network. Module statistics of a reference network (human) can be used to quantify which aspects, termed “patterns of connectivity,” are preserved in a second test (mouse) network^47^. We found that module eigengenes and many of the preservation statistics calculated between the human and mouse networks were conserved (Fig. S3). Overall, the patterns of connectivity among the blue, turquoise, and red modules in the human network were well-preserved in the mouse network, providing strong evidence that the human network is driven largely by pathogenic tau and is conserved across disease states (Fig. S4).

### Brains of tau transgenic *Drosophila* exhibit canonical cellular hallmarks of EMT

Based on identification of modules associated with EMT, cancer, growth and morphology in our human and mouse network analyses, we turned to *Drosophila* to investigate a potential link between pathogenic forms of tau and EMT-like phenotypes. Panneuronal expression of human wild-type tau and disease-associated tau mutants in *Drosophila* recapitulates many aspects of Alzheimer’s disease and related primary tauopathies including progressive neurodegeneration^48^, DNA damage^49^, and synapse loss^50^. In addition, neurons of tau transgenic *Drosophila* undergo an abortive cell cycle activation via a neurodegenerative process that shares many features of metastatic cancer cells, including over-stabilization of filamentous actin^11,51^, nuclear pleomorphism^14,52^, loss of heterochromatin-mediated transcriptional silencing^16,17^, and activation of transposable elements^53–55^. We analyzed canonical features of EMT in a *Drosophila* model of tauopathy that features panneuronal expression of a disease-associated mutant form of tau (tau^R406W^, referred to hereafter as “tau transgenic *Drosophila*” for simplicity)^48^. We performed all analyses in ten-day-old adults unless otherwise noted, as this well-described model exhibits a moderate degree of toxicity at day ten of adulthood that is well-suited for genetic analyses.

During EMT, changes in the actin cytoskeleton cause downregulation of adhesion molecules such as cadherin 1 and catenin alpha 1, and upregulation of cadherin 2^56,57^. We found that shotgun and α-catenin, the *Drosophila* homologs of human cadherin 1 and catenin alpha 1, were significantly decreased in the brains of tau transgenic *Drosophila* compared to controls (Fig. 3A-D). Immunostaining of Cadherin-N (CadN), the *Drosophila* homolog of human cadherin 2, revealed focal increases in CadN in brains of tau transgenic *Drosophila* (Fig. 3E), while total protein levels of CadN were significantly decreased in heads of tau transgenic *Drosophila* (Fig. 3F, Fig. S5A). As neurons are clearly not epithelial cells, we also investigated cellular adhesion proteins that are important for neuron-specific functions in *Drosophila*. We detected a significant reduction in Neuroglian (Nrg) and Fasciclin 2 (Fas2) (Fig. S5B-D), which regulate synapse formation, axon pathfinding, and neurite extension^58–60^, indicating that cellular adhesion proteins that are important for neuronal function are also depleted in tau transgenic *Drosophila.* Taken together, depletion of adhesion molecules that are canonical hallmarks of EMT, along with focal elevation of CadN in brains of tau transgenic *Drosophila,* suggest that pathogenic tau drives neuronal changes that mimic the cellular phenotypes of EMT.

**Figure 3.**
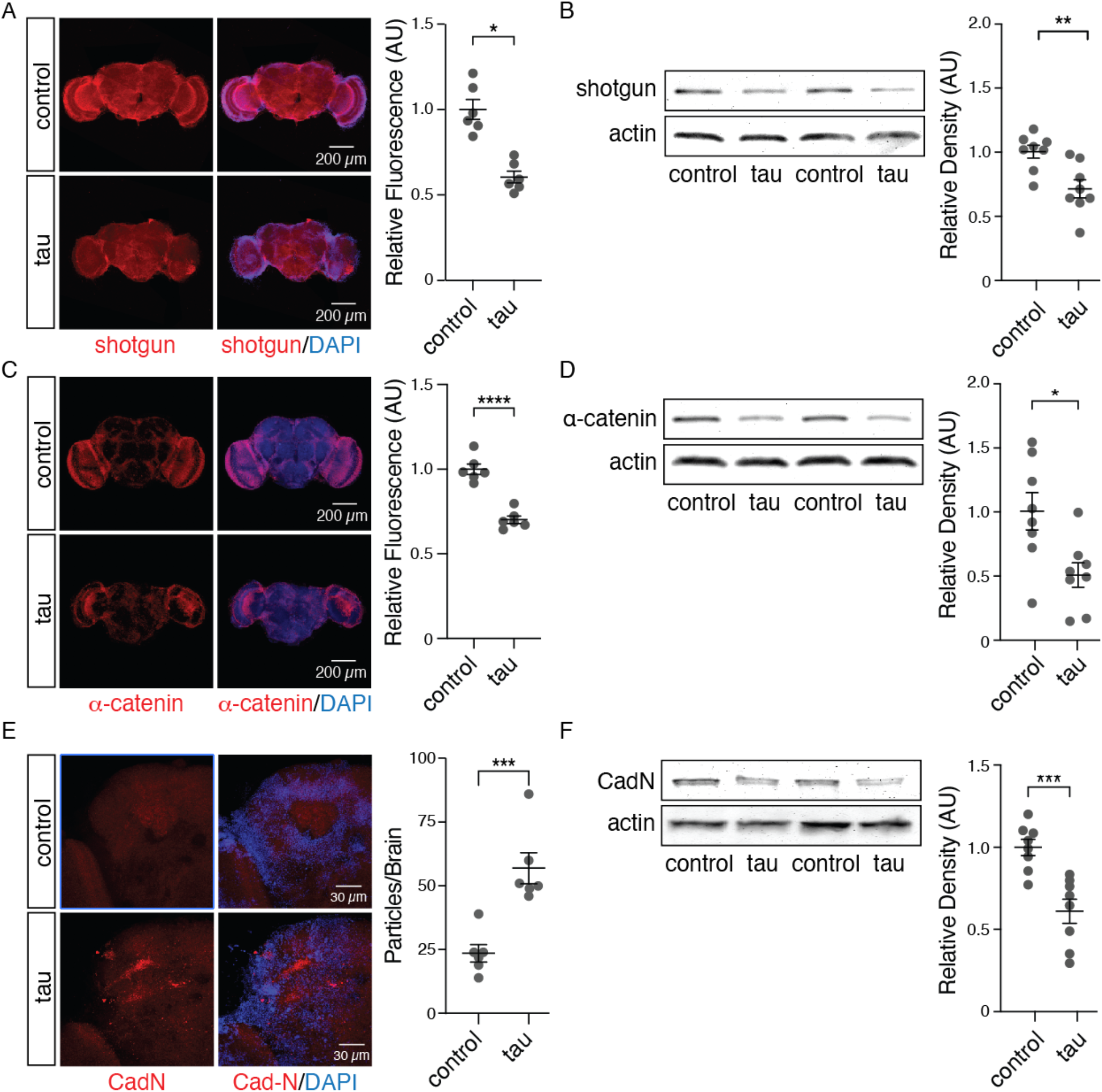
Hallmarks of EMT are conserved in brains of tau transgenic *Drosophila*. Decreased protein levels of shotgun **(A, B)** and α-catenin **(C, D)** in brains of tau transgenic *Drosophila* compared to control based on immunostaining and Western blotting (unpaired, two-tailed Student’s t-test). **(E)** Increased focal CadN by immunostaining while **(F)** total levels of CadN are decreased in tau transgenic *Drosophila* by Western blotting (unpaired, two-tailed Student’s t-test). n = 6-8 biologically independent replicates per genotype. All flies were ten days old. Values are mean ± s.e.m. *P < 0.05, **P < 0.005, ***P < 5.0×10^−4^, ****P < 5.0×10^−5^. Full genotypes are listed in Supplementary Table 3.

### Moesin activation drives filamentous actin formation and abortive cell cycle re-entry in brains of tau transgenic *Drosophila*

In humans, the Ezrin, Radixin and Moesin (ERM) proteins crosslink filamentous actin to the plasma membrane^61,62^. Aberrant activation of Moesin causes over-stabilization of the actin cytoskeleton, which mediates EMT and metastasis in many different cancers^63,64^. Filamentous actin is also over-stabilized in multiple models of tauopathy^65,66^, and mediates tau-induced abortive cell cycle activation^11^. Based on the presence of *Moesin* as a hub gene across human and mouse tauopathy networks, the ability of Moesin activation to mediate EMT through its effects on actin dynamics, and EMT-like cellular phenotypes in brains of tau transgenic *Drosophila,* we hypothesized that Moesin activation drives over-stabilization of filamentous actin and subsequent abortive cell cycle activation in tauopathy.

We found that panneuronal overexpression of a constitutively active form of Moesin, Moesin^T559D^ (hereafter referred to as “Moesin^CA^”), was sufficient to significantly elevate levels of filamentous actin in the adult *Drosophila* brain based on phalloidin staining (Fig. 4A). Conversely, panneuronal RNAi-mediated knockdown of Moesin in tau transgenic *Drosophila* significantly decreased levels of filamentous actin (Fig. 4B). Consistent with links between Moesin activation, filamentous actin, and cell cycle activation, we found that focal increases in Moesin occured in cell populations enriched for filamentous actin (Fig. 4C) and neurons that have aberrantly activated the cell cycle (Fig. 4D) in brains of tau transgenic *Drosophila*. In addition, panneuronal RNAi-mediated knockdown of Moesin significantly suppressed, while overexpression of constitutively active Moesin significantly enhanced, abortive cell cycle activation in brains of tau transgenic *Drosophila* (Fig. 4E, F). Collectively, these data suggest that aberrant Moesin activation mediates actin overstabilization and aberrant cell cycle activation in tauopathy.

**Figure 4.**
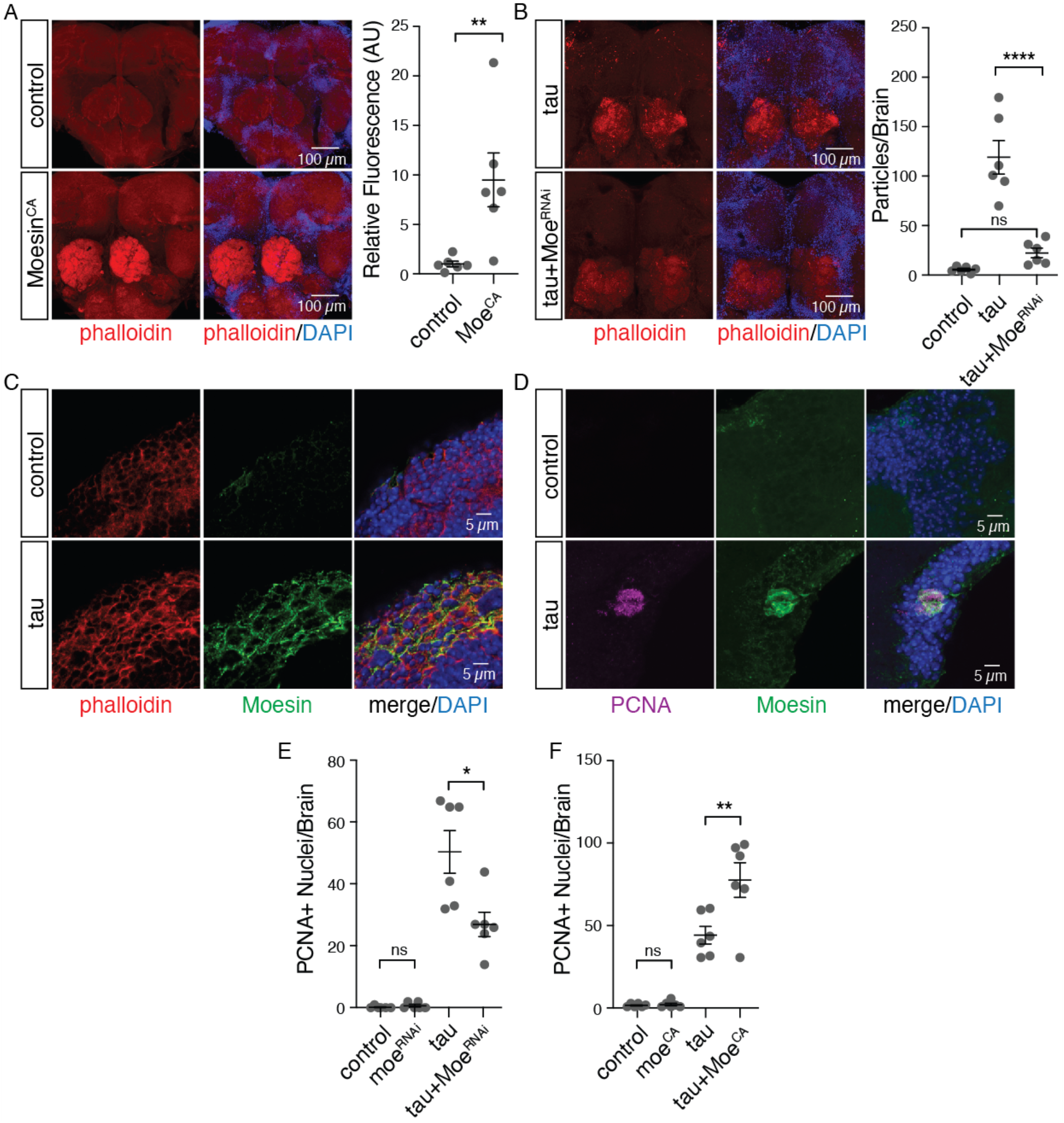
Moesin activation drives filamentous actin formation and abortive cell cycle activation in brains of tau transgenic *Drosophila.* **(A)** Increased fluorescence of phalloidin in the central brain of *Drosophila* harboring a constitutively active *Moesin* mutant relative to control (unpaired, two-tailed Student’s t-test). **(B)** RNAi-mediated Moesin knockdown decreased high signal foci phalloidin particles in brains of tau transgenic *Drosophila* (one-way ANOVA, Tukey’s test). **(C)** Moesin and phalloidin coincided in the medulla in control and tau transgenic *Drosophila* brains. **(D)** PCNA and Moesin colocalized in all observations of positive PCNA staining in tau transgenic *Drosophila*. PCNA staining indicated that Moesin knockdown significantly suppressed tau-induced cell cycle activation **(E)**, while constitutive activation of Moesin significantly enhanced tau-induced cell cycle activation **(F)**(one-way ANOVA, Tukey’s test). All flies were ten days old. Values are mean ± s.e.m., n = 6 biologically independent replicates per genotype, *P<0.05, **P < 5.0×10^−3^, ****P < 5.0×10^−5^. Full genotypes are listed in Supplementary Table 3.

### Pathogenic tau disrupts the cellular program that maintains terminal neuronal differentiation

As the EMT is a process involving loss of cellular identity, and canonical hallmarks of EMT are present in tau transgenic *Drosophila*, we next investigated the effects of tau on the cellular program that maintains terminal differentiation in *Drosophila* neurons. Using RNA-seq data generated from heads of control and tau transgenic *Drosophila*^67^, co-expression analysis identified a module (turquoise) that is strongly associated with biological processes such as neuronal developement, cellular growth, and RNA processing (Fig. S6A, Table S4), similar to our findings in human and mouse network analyses. Within the turquoise module, we identify *prospero,* which encodes a terminal neuronal selector protein that promotes and maintains neuronal differentiation in *Drosophila*^68^, as a hub gene. Prospero acts as a transcription factor to promote and maintain neuronal differentiation by silencing genes associated with development and activating genes that commit cells to the neuronal lineage^68^. A hypergeometric test indicated that genes known to be transcriptionally regulated by prospero^68^ are over-represented in the network (1.29 fold enrichment, p=1.15×10^−8^), and 23 of the 148 prospero-regulated genes were differentially expressed in tau transgenic *Drosophila* (Fig. S6B).

Based on immunostaining and western blotting, we found that prospero protein levels are significantly reduced in brains of tau transgenic *Drosophila* (Fig. 5A, B). To determine if prospero depletion is mediated by Moesin- and EMT-related processes, we genetically manipulated filamentous actin and Moesin in *Drosophila* neurons in the absence of transgenic tau. We found that over-stabilization of filamentous actin via overexpression of the actin polymerizing protein, Wiskott-Aldrich Syndrome protein (WASp)^69^ was sufficient to deplete prospero at the protein level (Fig. 5C, D), as was constitutively active Moesin (Fig. 5E, F).

**Figure 5.**
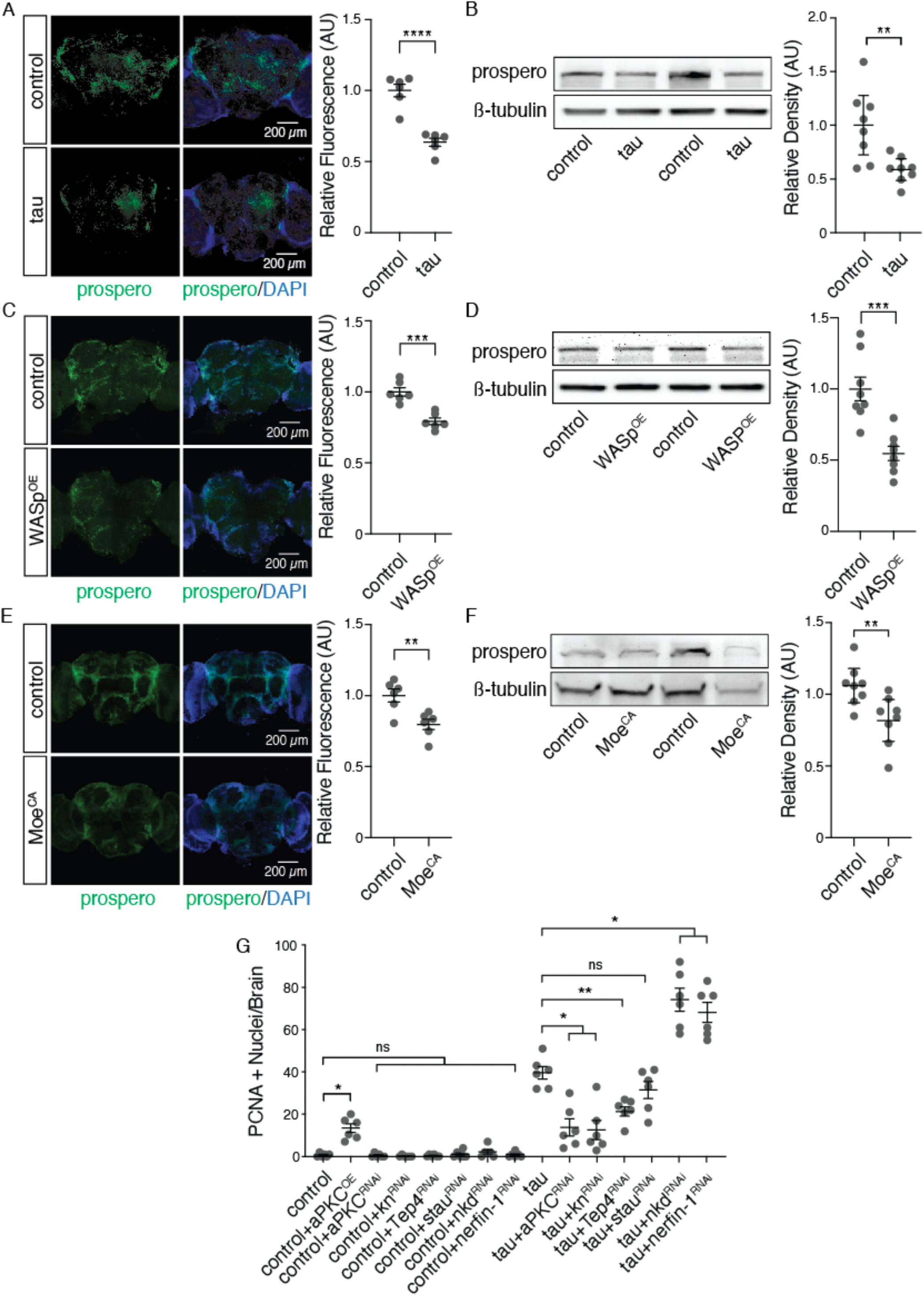
Pathogenic tau promotes abortive cell cycle activation by inducing the loss of terminal neuronal differentiation. Immunofluorescent staining and western blot analysis show decreased prospero protein levels in tau transgenic **(A, B)**, WASpOE **(C, D)** and MoesinCA **(E, F)** *Drosophila* (unpaired, two-tailed Student’s t-test). **(G)** Genetic manipulation of transcriptional targets of prospero suppressed tau-induced cell cycle activation based on PCNA staining (one-way ANOVA, Tukey’s test). n = 6-8 biologically independent replicates per genotype. All flies were ten days old. Full genotypes are listed in Supplementary Table 3. Values are mean ± s.e.m. *P < 0.05, **P < 0.005, ***P < 5.0×10^−4^.

We next asked if prospero depletion causally mediates tau-induced cell cycle activation. We found that both panneuronal overexpression or knockdown of prospero was toxic in both control and tau transgenic *Drosophila*, which is unsurprising given the central role of prospero in neurodevelopment. We instead asked if genetic manipulation of transcriptional targets of prospero modified tau-induced cell cycle activation. Consistent with a causal association between tau-induced abortive cell cycle activation and the cellular program regulated by prospero, we found that genetic manipulation of transcriptional targets of prospero significantly modified tau-induced activation of the cell cycle (Fig. 5G). Taken together, these data suggest that pathogenic tau drives abortive cell cycle activation by disrupting the cellular program that maintains terminal differentiation.

## Discussion

Since the initial discovery in 1996 linking pathogenic forms of tau to upregulation of the cell cycle-related protein p16 in neurons of the adult Alzheimer’s disease brain^70^, a wealth of literature has implicated tau as a driver of abortive cell cycle activation in neurons^71–74^. Work in multiple model systems has identified a series of cellular events connecting pathogenic forms of tau to cell cycle re-entry, including over-stabilization of the cytoskeleton^75,76^, disruption of nuclear architecture^77,78^, loss of heterochromatin-mediated gene silencing^79,80^, and activation transposable elements^53,55^. Each of these cellular phenotypes induced by tau are also features of cancerous cells^15,81–83^. The parallels between cancer, in which cells lose cellular identity, and tauopathy inspired our initial investigation into the effects of tau on cellular programs that maintain a terminally differentiated state.

Our network analysis of postmortem brain tissue from patients with Alzheimer’s disease and tau transgenic mice at three stages of disease identified *Moesin* as a hub gene within an expression module associated with cancer, EMT, growth and development. Moving into the *Drosophila* brain for mechanistic studies, we found that features of EMT are conserved in *Drosophila* tauopathy, as well as a causal association between Moesin activation, filamentous actin formation, and cell cycle re-entry. Taken together, these findings suggest that the cancer/EMT-related expression module in late-stage human Alzheimer’s disease brains is a consequence of pathogenic forms of tau and is conserved throughout disease stage. Our findings converge with those of the National Institute on Aging’s Accelerating Medicines Partnership – Alzheimer’s Disease consortium, which has recently nominated Moesin as a drug target for Alzheimer’s disease (https://agora.ampadportal.org/genes/(genes-router:gene-details/ENSG00000147065) based on genomic and proteomic data from human Alzheimer’s disease samples. Our identification of *Moesin* as a hub gene in human and mouse tauopathy networks aligns with the findings of the consortia, and our studies in *Drosophila* provide the mechanistic insight into the consequences of Moesin activation in tauopathy that are critical for drug development.

Having discovered that pathogenic tau drives cellular phenotypes that are canonical features of EMT, we next explored the effects of pathogenic tau on the cellular program that maintains terminal neuronal differentiation in *Drosophila.* We found that prospero, a key transcription factor that activates genes associated with neuronal differentiation and silences genes associated with neuronal development^68^ was depleted in tau transgenic *Drosophila,* and is likely downstream of aberrant tau-induced Moesin activation and consequent over-stabilization of filamentous actin. Our finding that transcriptional targets of prospero significantly modify tau-induced cell cycle activation suggests a causal relationship between tau, the cellular program that maintains terminal differentiation, and abortive cell cycle re-entry. While not tested in the current study, we speculate that nuclear architecture disruption^14^, heterochromatin decondensation^16^, and retrotransposon activation^53^ in tauopathy are a consequence of disrupted neuronal identity.

Identifying disease mechanisms that are causally associated with neurotoxicity and conserved across disease stage is critical to overcome the challenge of treating neurodegenerative disorders that span several decades. By combining bioinformatic approaches in human and mouse tauopathy with mechanistic studies in a *Drosophila* model of tauopathy, we identify Moesin as a key player connecting pathogenic forms of tau to loss of neuronal identity. Our results provide novel insights into the connection between tau and aberrant cell cycle activation and identify new targets for therapeutic intervention.

## Methods

### Human, mouse, and *Drosophila* model systems

#### Human

RNA-seq data was available for 76 patients with Alzheimer’s disease (42.1% male, 57.9% female) and 48 non-demented controls (52.1% males, 47.9% female). Covariate information including sex, age at death, RNA integrity number and sequencing flow-cell are available in the AMP-AD Knowledge Portal (Synapse ID: syn3163039).

#### Mice

Female rTg4510 and non-transgenic control mice were obtained from the Jackson Laboratory. Mice were group-housed at 22 °C with a 12 h light/dark cycle with food and water available *ad libitum*. 17 tau transgenic mice and 18 non-transgenic mice aged to three, six, and nine months. After anesthesia with 2% isoflurane, cardiac perfusion was performed using 2X PhosSTOP phosphatase inhibitors (Roche, Indianapolis, IN, USA) and 1X complete protease inhibitors (Roche) in PBS (Thermo Fisher Scientific, Waltham, MA, USA). Brains were then removed, and hemispheres were separated at the midline. The right hemisphere was further dissected to isolate the cerebellum, brainstem, hippocampus, and remaining forebrain and snap frozen. Hippocampus was used for RNA-seq. All experimental procedures were performed according to the National Institute of Health Guidelines for the Care and Use of Laboratory Animals and were approved by the Institutional Animal Care and Use Committee of the University of Texas MD Anderson Cancer Center.

#### Drosophila

Crosses and aging were performed at 25 °C with a 12 h light/dark cycle at 60% relative humidity on a standard diet (Bloomington formulation). Full genotypes are listed in Supplementary Table 3. Panneuronal expression of transgenes, including RNAi-mediated knockdown, in *Drosophila* was achieved using the GAL4/UAS system with the *elav* promoter driving GAL4 expression^84^. The *elav-GAL4* line was obtained from the Bloomington Stock Center (BDSC 458). The *UAS-tau*^*R406W*^ line was a gift from Dr. Mel Feany. An equal number of males and females were used in all *Drosophila* assays.

### RNA sequencing and differential gene expression analyses

#### Human

Whole-transcriptome data was downloaded from the AMP-AD Knowledge Portal (Synapse ID: syn3163039). Gene expression data from temporal cortex was generated by the Mayo Clinic Brain Bank using Illumina HiSeq 2000-based next-generation 101 bp paired-end sequencing. FASTQ files were trimmed with Trimmomatic (v.0.36)^85^ to remove adapters and low-quality reads. FastQC^86^ was used to evaluate read quality before and after trimming. Trimmed FASTQ files were mapped and aligned to the *Homo sapiens* transcriptome (Gencode v31) using Salmon (v.0.13.1)^87^. Differential expression analysis was performed using DESeq2 (v1.24)^88^. Trimmomatic and Salmon tools were run using the resources provided by the UT Health SA Bioinformatics Core Facility. Genes with an adjusted p value of less than 0.05 were considered significant.

#### Mouse

Hippocampal RNA was isolated using Qiagen RNeasy Mini kit and Qiagen TissueRuptor (Qiagen, Germantown, MD, USA) per the manufacturer’s recommendations. RNA-Seq was performed using Illumina HiSeq 4000 with 76 bp paired-end reads. FASTQ files were trimmed with Trimmomatic (v.0.36)^85^ to remove adapters and low-quality reads. FastQC^86^ was used to evaluate the quality of the reads before and after trimming. Trimmed FASTQ files were mapped and aligned to the *Mus musculus* transcriptome (Gencode M22) using Salmon (v.0.13.1)^87^. Differential expression analysis was performed using DESeq2 (v1.24)^88^. Mouse Trimmomatic and Salmon tools were run using resources on the TACC Lonestar5 cluster. Genes with an adjusted p value of less than 0.05 were considered significant.

#### Drosophila

RNA-seq data from control and tau transgenic *Drosophila*^67^ were obtained from the Gene Expression Omnibus (GEO) database (GSE152278).

### Weighted gene co-expression network analyses

Each of the networks in this study were constructed using the R WGCNA package and methodologies previously described^33^. Here, gene expression data was normalized by transcripts per million (TPM) and log base two transformation. From more than 15,000 genes, 8,000 of the most varying genes were preliminarily selected for network construction. Genes were removed if they contained too many missing values (minimal fraction = 1/2), if mean expression was less than two TPM, if they had zero variance, or if they were not conserved in humans. Outlier samples were detected by hierarchical clustering using the R core Stats package. In order to obtain biologically meaningful networks and understand the directionality of node profiles, we constructed signed hybrid adjacency matrices where the absolute value of the Pearson correlation measures gene is the co-expression similarity, and aij represents the resulting adjacency that measures the connection strengths *a*_*_ij_*_ = *|cor(X*_*i*_, *X*_*j*_*)|* ^*ß*^. Network connectivity 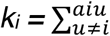 is defined as the sum of connection strengths with other genes. Soft-thresholding powers (ß) were selected using the scale-free criterion in which the network connectivity distribution of nodes approximately followed inverse power law *p(k)~k*^*~γ*89^. Due to limitations in data visualization software, networks were further restricted to the 5,000 most connected genes. Modules were defined as genes with high topological overlap where the overlap between genes *i* and *j* was measured using 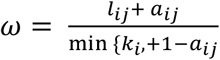. Modules were identified by average linking hierarchical clustering along with the distance calculated from the topological overlap matrix as a measure of dissimilarity 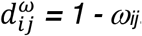. Module cut heights ranged from 0.1-0.25 based on the number of modules detected and cluster distancing. Only co-expressed genes in groups of 100 genes or more were considered modules. Hub genes for each module were identified by ranking genes according to their intramodular connectivity (k_in_) and selecting the top 1-5% of the most connected genes. In each case, modules were assessed for enrichment in biological processes using the enrichGO algorithm provided by clusterProfiler (v3.04)^90^. Associations with biological processes were considered significant if adjusted p values (false discovery rate) were less than 0.05. To identify gene-disease associations for each module, we utilized DOSE (v3.11)^91^ in conjunction with the enrichDGN algorithm. Gene-disease associations were considered significant if adjusted p values (false discovery rate) were less than 0.05.

#### Module preservation analysis

Module preservation analysis was performed using methodologies previously described^47^. The gene clustering dendrogram of the mouse network was re-created using the same network construction techniques as in the human network. To restrict our analysis to the most preserved and connected genes, we only included genes with scaled connectivities greater than 0.1, resulting in 3,597 genes. Determination of preservation statistics was performed using the modulePreservation function from the WGCNA package and corrected for multiple testing using Bonferroni’s correction. A comprehensive set of module preservation statistics is provided in Figs. S3-4. See Langfelder *et al*. for complete list of definitions and glossary^47^.

#### Principal component analyses

For multi-dimensional scaling plots depicted in Figs. 1 and 2, module eigengene and whole-network matrices were analyzed using the prcomp function from the R core package Stats. Whole networks and module eigengenes from each of their respective networks were analyzed using the measure of dissimilarity previously calculated.

#### Prospero target analysis

A hypergeometric test (R core package Stats) was used to determine enrichment of prospero targets^68^ in the list of genes used in WGCNA (Supplementary Table 5) for *Drosophila*. Briefly, fold enrichment of prospero target genes was performed as previously computed by DAVID^92^ where 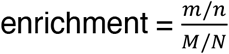. For background, we utilized the 8,000 most varying genes from the human expression data. In total, 148 prospero target genes were identified from the 5,000 genes that made up the *Drosophila* network.

### Immunofluorescence and histology

For prospero, CadN, α-catenin, shotgun, Nrg, and Fas2 immunofluorescence, *Drosophila* brains were dissected in PBS, fixed in methanol for 10 min and adhered to microscope slides. Slides were rinsed in diH20 and washed using 1X PBS followed by blocking with 2% milk in PBS plus 0.3% TritonX (PBS_Tr_) for 30 min. Slides were incubated with primary antibodies diluted in blocking solution overnight at 4 °C. After incubation with primary antibodies, slides were washed with 0.3% PBS_Tr_ and incubated with Alexa488-, Alexa555-, or Alexa647-conjugated secondary antibodies diluted in blocking solution for 2 h at room temperature. Lastly, slides were washed again and incubated with DAPI for 2 min to stain nuclei before cover slipping with fluorescent mounting medium. For phalloidin staining, *Drosophila* brains were fixed in 4% PFA for 10 min and prepared for staining according to the manufacturer’s protocol (Cell Signaling Technology). Brains were visualized by confocal microscopy (Zeiss LSM 780 NLO with Examiner, Zeiss LSM 810 with Airyscan), and ImageJ^93^ was used for analysis. PCNA staining was performed using 4 μm sections from formalin-fixed, paraffin-embedded *Drosophila* heads. Sections were adhered to microscope slides then deparaffinized and dehydrated using a xylene and ethanol series of rinses and washes. To improve signal detection, slides were heated to 100°C for 15 min in 1L of 10mM sodium citrate in 0.05% Tween 20, pH 6.0. Slides were then washed in PBS and blocked using 2% milk in 0.3% PBS_Tr_ for 30 min. Next, slides were incubated with PCNA antibodies diluted in blocking solution overnight at 4°C. Secondary detection was performed with Diaminobenzidine according to the manufacturer’s protocol (Vector Laboratories). PCNA-positive neurons were counted throughout the entire brain by bright field microscopy (Nikon Eclipse Ci-L). Antibodies, reagents, concentrations, and sources are listed in Supplementary Table 6.

#### Particle Analysis

To quantify high signal foci from images of CadN and phalloidin staining in *Drosophila* brains, we utilized the Analyze Particles tool from ImageJ^93^. Briefly, stacked images were converted to z-projections using MaxEntropy thresholding to exclude low signal and background. Z-projected images were then converted to 8-bit binary images using MaxEntropy thresholding. To reliably quantify the number of particles per brain we excluded particles outside of the brain, with size less than 0.1 pixel^2 or greater than 100 pixels^2. Circularity was left to the default setting.

### Western blotting

Frozen *Drosophila* heads were homogenized in 15 μl of 2X Laemmli sample buffer, heated for 10 min at 70°C, and analyzed by 4–20% or 7.5% SDS–PAGE using the Bio-Rad mini-PROTEAN Tetra Cell. Polyacrylamide gels were transferred at 4°C for 90 min at 90v to nitrocellulose or PVDF membranes using the Bio-Rad Mini Trans-Blot Cell and Towbin Buffer^94^. Equal loading was assessed by Ponceau S staining. Membranes were then incubated at 4°C for 30 min in a blocking solution made up of 2% milk in PBS plus 0.05% Tween (PBS_Tw_) followed by incubation with primary antibodies overnight at 4°C with gentle rocking. Membranes were then washed using 0.05% PBS_Tw_ and incubated with HRP-conjugated secondary antibodies for 2 h at room temperature. Blots were developed with an enhanced chemiluminescent substrate and imaged using the ProteinSimple FluorChem HD2 system. Antibodies, reagents, concentrations, and sources are listed in Supplementary Table 6. Full scans of western blots are provided in Figure S7.

### Statistical analyses

Every reported *n* is the number of biologically independent replicates. Except when noted otherwise, statistical analyses were performed using a one-way ANOVA with Tukey test when comparing among multiple genotypes and a two-tailed, unpaired Student’s t-test when comparing two genotypes. Data distribution was assumed to be normal, but this was not formally tested. For RNA-seq analysis, a two-sided Wald test was used to calculate false discovery rates (FDR-adjusted p value)^95^. A p value less than 0.05 was considered significant unless otherwise specified. Sample sizes are similar to or greater than those reported in previous publications. Samples were randomized in all *Drosophila* studies. Investigators were blinded to genotype in all immunohistochemistry and immunofluorescence. Full statistical analyses are provided in Supplementary Text 1. To improve transparency and increase reproducibility, detailed information on experimental design and reagents can be accessed in the Reporting Summary.

## Acknowledgements

Images were generated in part at the Core Optical Imaging Facility, which is supported by UTHSCSA, NIH-NCI P30 CA54174 (CTRC at UTHSCSA) and NIH-NIA P01AG19316. Special thanks to the Kiehart lab and Dr. Janice Crawford for providing critical aliquots of Moesin antibodies, and to Dr. Mel Feany for providing the *UAS-tau*^*R406W*^ *Drosophila* stock. This study was supported by the National Institute on Aging (P30 AG13319) and the Briscoe Women’s Health Scholar Fund. AB was supported by T32 AG021890 and F31 NS108657. PR was supported by R25 GM095480. The Neurodegeneration Consortium (JR) is supported by the Robert A and Renee E Belfer Foundation, the Oskar Fisher Project, and other philanthropic sources. BF and MG were supported by R01 AG057896. Stocks obtained from the Bloomington *Drosophila* Stock Center (NIH P40 OD018537) were used in this study. The Mayo human RNAseq study data was led by N. Ertekin-Taner (Mayo Clinic) as part of the multi-PI U01 AG046139 (MPIs Golde, Ertekin-Taner, Younkin, Price) using samples from the following source:

▪ The Mayo Clinic Brain Bank. Data collection was supported through funding by NIA grants P50 AG016574, R01 AG032990, U01 AG046139, R01 AG018023, U01 AG006576, U01 AG006786, R01 AG025711, R01 AG017216, R01 AG003949, NINDS grant R01 NS080820, CurePSP Foundation, and support from Mayo Foundation.

## Notes

### Competing Interest Statement

The authors have declared no competing interest.

## Literature Cited

1. Vissers, J. H. A. et al. The Scalloped and Nerfin-1 Transcription Factors Cooperate to Maintain Neuronal Cell Fate. Cell Rep. 25, 1561–1576.e7 (2018).

2. Yoshikawa, K. Cell cycle regulators in neural stem cells and postmitotic neurons. Neuroscience Research vol. 37 1–14 (2000).

3. Ajioka, I. Coordination of proliferation and neuronal differentiation by the retinoblastoma protein family. Development Growth and Differentiation vol. 56 324–334 (2014).

4. Arendt, T., Holzer, M. & Gärtner, U. Neuronal expression of cycline dependent kinase inhibitors of the INK4 family in Alzheimer’s disease. J. Neural Transm. 105, 949–960 (1998).

5. Busser, J., Geldmacher, D. S. & Herrup, K. Ectopic Cell Cycle Proteins Predict the Sites of Neuronal Cell Death in Alzheimer’s Disease Brain. J. Neurosci. 18, 2801–2807 (1998).

6. Yang, Y., Mufson, E. J. & Herrup, K. Neuronal Cell Death Is Preceded by Cell Cycle Events at All Stages of Alzheimer’s Disease. J. Neurosci. 23, 2557–2563 (2003).

7. Liu, D. X. & Greene, L. A. Neuronal apoptosis at the G1/S cell cycle checkpoint. Cell Tissue Res. 305, 217–228 (2001).

8. Yang, Y., Geldmacher, D. S. & Herrup, K. DNA replication precedes neuronal cell death in Alzheimer’s disease. J. Neurosci. 21, 2661–2668 (2001).

9. Arendt, T. Cell Cycle Activation and Aneuploid Neurons in Alzheimer’s Disease. Mol. Neurobiol. 46, 125–135 (2012).

10. DuBoff, B., Götz, J. & Feany, M. B. Tau Promotes Neurodegeneration via DRP1 Mislocalization In Vivo. Neuron 75, 618–632 (2012).

11. Fulga, T. A. et al. Abnormal bundling and accumulation of F-actin mediates tau-induced neuronal degeneration in vivo. Nat. Cell Biol. 9, 139–148 (2007).

12. Li, Q. S., Lee, G. Y. H., Ong, C. N. & Lim, C. T. AFM indentation study of breast cancer cells. Biochem. Biophys. Res. Commun. 374, 609–613 (2008).

13. Clucas, J. & Valderrama, F. ERM proteins in cancer progression. J. Cell Sci. 128, 1253–1253 (2015).

14. Frost, B., Bardai, F. H. & Feany, M. B. Lamin Dysfunction Mediates Neurodegeneration in Tauopathies. Curr. Biol. 26, 129–136 (2016).

15. Zink, D., Fischer, A. H. & Nickerson, J. A. Nuclear structure in cancer cells. Nat. Rev. Cancer 4, 677–687 (2004).

16. Frost, B., Hemberg, M., Lewis, J. & Feany, M. B. Tau promotes neurodegeneration through global chromatin relaxation. Nat. Neurosci. 17, 357–366 (2014).

17. Zhu, Q. et al. BRCA1 tumour suppression occurs via heterochromatin-mediated silencing. Nature 477, 179–184 (2011).

18. Johnson, N. et al. Actin-filled nuclear invaginations indicate degree of cell de-differentiation. Differentiation 71, 414–424 (2003).

19. Talamas, J. A. & Capelson, M. Nuclear envelope and genome interactions in cell fate. Front. Genet. 6, 1–16 (2015).

20. Hsieh, J. & Gage, F. H. Epigenetic control of neural stem cell fate. Curr. Opin. Genet. Dev. 14, 461–469 (2004).

21. Weber, C. E., Li, N. Y., Wai, P. Y. & Kuo, P. C. Epithelial-Mesenchymal Transition, TGF-β, and Osteopontin in Wound Healing and Tissue Remodeling After Injury. J. Burn Care Res. 33, 311–318 (2012).

22. Yan, C. et al. Epithelial to Mesenchymal Transition in Human Skin Wound Healing Is Induced by Tumor Necrosis Factor-α through Bone Morphogenic Protein-2. Am. J. Pathol. 176, 2247–2258 (2010).

23. Hardy, K. M., Booth, B. W., Hendrix, M. J. C., Salomon, D. S. & Strizzi, L. ErbB/EGF Signaling and EMT in Mammary Development and Breast Cancer. J. Mammary Gland Biol. Neoplasia 15, 191–199 (2010).

24. Shankar, J. et al. Pseudopodial Actin Dynamics Control Epithelial-Mesenchymal Transition in Metastatic Cancer Cells. Cancer Res. 70, 3780–3790 (2010).

25. Shankar, J. & Nabi, I. R. Actin Cytoskeleton Regulation of Epithelial Mesenchymal Transition in Metastatic Cancer Cells. PLoS One 10, e0119954 (2015).

26. Haynes, J., Srivastava, J., Madson, N., Wittmann, T. & Barber, D. L. Dynamic actin remodeling during epithelial–mesenchymal transition depends on increased moesin expression. Mol. Biol. Cell 22, 4750–4764 (2011).

27. Lu, P., Weaver, V. M. & Werb, Z. The extracellular matrix: A dynamic niche in cancer progression. J. Cell Biol. 196, 395–406 (2012).

28. Moussalli, M. J. et al. Mechanistic contribution of ubiquitous 15-lipoxygenase-1 expression loss in cancer cells to terminal cell differentiation evasion. Cancer Prev. Res. 4, 1961–1972 (2011).

29. Ma, W. et al. Cell-Extracellular Matrix Interactions Regulate Neural Differentiation of Human Embryonic Stem Cells. BMC Dev. Biol. 8, 90 (2008).

30. Bonneh-Barkay, D. & Wiley, C. A. Brain Extracellular Matrix in Neurodegeneration. Brain Pathol. 19, 573–585 (2009).

31. Smith, L. R., Cho, S. & Discher, D. E. Stem Cell Differentiation is Regulated by Extracellular Matrix Mechanics. Physiology 33, 16–25 (2018).

32. Hobert, O. Regulation of Terminal Differentiation Programs in the Nervous System. Annu. Rev. Cell Dev. Biol. 27, 681–696 (2011).

33. Langfelder, P. & Horvath, S. WGCNA: An R package for weighted correlation network analysis. BMC Bioinformatics 9, 559 (2008).

34. Ashburner, M. et al. Gene Ontology: tool for the unification of biology. Nat. Genet. 25, 25–29 (2000).

35. Carbon, S. et al. The Gene Ontology Resource: 20 years and still GOing strong. Nucleic Acids Res. 47, D330–D338 (2019).

36. Piñero, J. et al. The DisGeNET knowledge platform for disease genomics: 2019 update. Nucleic Acids Res. 48, D845–D855 (2020).

37. Bassat, E. et al. The extracellular matrix protein agrin promotes heart regeneration in mice. Nature 547, 179–184 (2017).

38. Fernandez-L, A. et al. YAP1 is amplified and up-regulated in hedgehog-associated medulloblastomas and mediates Sonic hedgehog-driven neural precursor proliferation. Genes Dev. 23, 2729–2741 (2009).

39. Bhat, K. P. L. et al. The transcriptional coactivator TAZ regulates mesenchymal differentiation in malignant glioma. Genes Dev. 25, 2594–2609 (2011).

40. Abiatari, I. et al. Moesin-dependent cytoskeleton remodelling is associated with an anaplastic phenotype of pancreatic cancer. J. Cell. Mol. Med. 14, 1166–1179 (2010).

41. Langfelder, P. & Horvath, S. Eigengene networks for studying the relationships between co-expression modules. BMC Syst. Biol. 1, 54 (2007).

42. Patel, T. & Hobert, O. Coordinated control of terminal differentiation and restriction of cellular plasticity. Elife 6, 1–26 (2017).

43. Rossor, M. N., Fox, N. C., Freeborough, P. A. & Harvey, R. J. Clinical Features of Sporadic and Familial Alzheimer’s Disease. Neurodegeneration 5, 393–397 (1996).

44. Fox, N. C., Freeborough, P. A. & Rossor, M. N. Visualisation and quantification of rates of atrophy in Alzheimer’s disease. Lancet 348, 94–97 (1996).

45. Mirra, S. S. et al. Tau Pathology in a Family with Dementia and a P301L Mutation in Tau. J. Neuropathol. Exp. Neurol. 58, 335–345 (1999).

46. Ramsden, M. Age-Dependent Neurofibrillary Tangle Formation, Neuron Loss, and Memory Impairment in a Mouse Model of Human Tauopathy (P301L). J. Neurosci. 25, 10637–10647 (2005).

47. Langfelder, P., Luo, R., Oldham, M. C. & Horvath, S. Is My Network Module Preserved and Reproducible? PLoS Comput. Biol. 7, e1001057 (2011).

48. Wittmann, C. W. et al. Tauopathy in Drosophila: Neurodegeneration without neurofibrillary tangles. Science (80−.). 293, 711–714 (2001).

49. Khurana, V. et al. A neuroprotective role for the DNA damage checkpoint in tauopathy. Aging Cell vol. 11 360–362 (2012).

50. Merlo, P. et al. P53 prevents neurodegeneration by regulating synaptic genes. Proc. Natl. Acad. Sci. U. S. A. 111, 18055–18060 (2014).

51. Tavares, S. et al. Actin stress fiber organization promotes cell stiffening and proliferation of pre-invasive breast cancer cells. Nat. Commun. 8, 15237 (2017).

52. Bussolati, G., Marchiò, C., Gaetano, L., Lupo, R. & Sapino, A. Pleomorphism of the nuclear envelope in breast cancer: A new approach to an old problem. J. Cell. Mol. Med. 12, 209–218 (2008).

53. Sun, W., Samimi, H., Gamez, M., Zare, H. & Frost, B. Pathogenic tau-induced piRNA depletion promotes neuronal death through transposable element dysregulation in neurodegenerative tauopathies. Nat. Neurosci. 21, 1038–1048 (2018).

54. Goudarzi, K. M. et al. Reduced expression of PROX1 transitions glioblastoma cells into a mesenchymal gene expression subtype. Cancer Res. 78, canres.0320.2018 (2018).

55. Guo, C. et al. Tau Activates Transposable Elements in Alzheimer’s Disease. Cell Rep. 23, 2874–2880 (2018).

56. Christofori, G. New signals from the invasive front. Nature vol. 441 444–450 (2006).

57. Grünert, S., Jechlinger, M. & Beug, H. Diverse cellular and molecular mechanisms contribute to epithelial plasticity and metastasis. Nat. Rev. Mol. Cell Biol. 4, 657–665 (2003).

58. Baines, R. A., Seugnet, L., Thompson, A., Salvaterra, P. M. & Bate, M. Regulation of synaptic connectivity: Levels of fasciclin II influence synaptic growth in the Drosophila CNS. J. Neurosci. 22, 6587–6595 (2002).

59. Zhang, H. et al. Endocytic Pathways Downregulate the L1-type Cell Adhesion Molecule Neuroglian to Promote Dendrite Pruning in Drosophila. Dev. Cell 30, 463–478 (2014).

60. Godenschwege, T. A., Kristiansen, L. V., Uthaman, S. B., Hortsch, M. & Murphey, R. K. A Conserved Role for Drosophila Neuroglian and Human L1-CAM in Central-Synapse Formation. Curr. Biol. 16, 12–23 (2006).

61. Bretscher, A., Edwards, K. & Fehon, R. G. ERM proteins and merlin: Integrators at the cell cortex. Nature Reviews Molecular Cell Biology vol. 3 586–599 (2002).

62. Hiruma, S., Kamasaki, T., Otomo, K., Nemoto, T. & Uehara, R. Dynamics and function of ERM proteins during cytokinesis in human cells. FEBS Letters vol. 591 3296–3309 (2017).

63. Sikorska, J., Gaweł, D., Domek, H., Rudzińska, M. & Czarnocka, B. Podoplanin (PDPN) affects the invasiveness of thyroid carcinoma cells by inducing ezrin, radixin and moesin (E/R/M) phosphorylation in association with matrix metalloproteinases 06 Biological Sciences 0601 Biochemistry and Cell Biology 06 Biological Scien. BMC Cancer 19, 1–17 (2019).

64. Bartova, M. et al. Expression of ezrin and moesin in primary breast carcinoma and matched lymph node metastases. Clin. Exp. Metastasis 34, 333–344 (2017).

65. Whiteman, I. T. et al. Activated Actin-Depolymerizing Factor/Cofilin Sequesters Phosphorylated Microtubule-Associated Protein during the Assembly of Alzheimer-Like Neuritic Cytoskeletal Striations. J. Neurosci. 29, 12994–13005 (2009).

66. Tracy, T. E. & Gan, L. Acetylated tau in Alzheimer’s disease: An instigator of synaptic dysfunction underlying memory loss. BioEssays 39, 1600224 (2017).

67. Mahoney, R. et al. Pathogenic Tau Causes a Toxic Depletion of Nuclear Calcium Mediated by BK Channels. SSRN Electron. J. (2020) doi:10.2139/ssrn.3519898.

68. Choksi, S. P. et al. Prospero Acts as a Binary Switch between Self-Renewal and Differentiation in Drosophila Neural Stem Cells. Dev. Cell 11, 775–789 (2006).

69. Berger, S. et al. WASP and SCAR have distinct roles in activating the Arp2/3 complex during myoblast fusion. J. Cell Sci. 121, 1303–1313 (2008).

70. Arendt, T., Rödel, L., Gärtner, U. & Holzer, M. Expression of the cyclin-dependent kinase inhibitor p16 in Alzheimer’s disease. Neuroreport 7, 3047–3049 (1996).

71. McShea, A. et al. Neuronal cell cycle re-entry mediates Alzheimer disease-type changes. Biochim. Biophys. Acta - Mol. Basis Dis. 1772, 467–472 (2007).

72. Seward, M. E. et al. Amyloid-β signals through tau to drive ectopic neuronal cell cycle re-entry in alzheimer’s disease. J. Cell Sci. 126, 1278–1286 (2013).

73. Jaworski, T. et al. AAV-tau mediates pyramidal neurodegeneration by cell-cycle re-entry without neurofibrillary tangle formation in wild-type mice. PLoS One 4, e7280 (2009).

74. Andorfer, C. et al. Cell-cycle reentry and cell death in transgenic mice expressing nonmutant human tau isoforms. J. Neurosci. 25, 5446–5454 (2005).

75. Torres-Cruz, F. M. et al. Expression of Tau Produces Aberrant Plasma Membrane Blebbing in Glial Cells Through RhoA-ROCK-Dependent F-Actin Remodeling. J. Alzheimer’s Dis. 52, 463–482 (2016).

76. Cabrales Fontela, Y. et al. Multivalent cross-linking of actin filaments and microtubules through the microtubule-associated protein Tau. Nat. Commun. 8, 1–12 (2017).

77. Fernández-Nogales, M. et al. Tau-positive nuclear indentations in P301S tauopathy mice. Brain Pathol. 27, 314–322 (2017).

78. Eftekharzadeh, B. et al. Tau Protein Disrupts Nucleocytoplasmic Transport in Alzheimer’s Disease. Neuron 99, 925–940.e7 (2018).

79. Mansuroglu, Z. et al. Loss of Tau protein affects the structure, transcription and repair of neuronal pericentromeric heterochromatin. Sci. Rep. 6, 33047 (2016).

80. Frost, B. Alzheimer’s disease: An acquired neurodegenerative laminopathy. Nucleus 7, 275–283 (2016).

81. Burns, K. H. Transposable elements in cancer. Nature Reviews Cancer vol. 17 415–424 (2017).

82. Dutta, P. et al. Unphosphorylated STAT3 in heterochromatin formation and tumor suppression in lung cancer. BMC Cancer 20, 145 (2020).

83. Valakh, V., Frey, E., Babetto, E., Walker, L. J. & DiAntonio, A. Cytoskeletal disruption activates the DLK/JNK pathway, which promotes axonal regeneration and mimics a preconditioning injury. Neurobiol. Dis. 77, 13–25 (2015).

84. Brand, A. H. & Perrimon, N. Targeted gene expression as a means of altering cell fates and generating dominant phenotypes. Development 118, 401–415 (1993).

85. Bolger, A. M., Lohse, M. & Usadel, B. Trimmomatic: A flexible trimmer for Illumina sequence data. Bioinformatics 30, 2114–2120 (2014).

86. S., A. FastQC: a quality control tool for high throughput sequence data. Babraham Bioinforma. (2010).

87. Patro, R., Duggal, G., Love, M. I., Irizarry, R. A. & Kingsford, C. Salmon provides fast and bias-aware quantification of transcript expression. Nat. Methods 14, 417–419 (2017).

88. Love, M. I., Huber, W. & Anders, S. Moderated estimation of fold change and dispersion for RNA-seq data with DESeq2. Genome Biol. 15, 1–21 (2014).

89. Zhang, B. & Horvath, S. A general framework for weighted gene co-expression network analysis. Stat. Appl. Genet. Mol. Biol. 4, 17 (2005).

90. Yu, G., Wang, L. G., Han, Y. & He, Q. Y. ClusterProfiler: An R package for comparing biological themes among gene clusters. Omi. A J. Integr. Biol. 16, 284–287 (2012).

91. Yu, G., Wang, L.-G., Yan, G.-R. & He, Q.-Y. DOSE: an R/Bioconductor package for Disease Ontology Semantic and Enrichment analysis. Bioinformatics 31, 608–609 (2015).

92. Jiao, X. et al. DAVID-WS: a stateful web service to facilitate gene/protein list analysis. Bioinformatics 28, 1805–1806 (2012).

93. Schneider, C. A., Rasband, W. S. & Eliceiri, K. W. NIH Image to ImageJ: 25 years of image analysis. Nature Methods vol. 9 671–675 (2012).

94. Towbin, H., Özbey, Ö. & Zingel, O. An immunoblotting method for high-resolution isoelectric focusing of protein isoforms on immobilized pH gradients. Electrophoresis 22, 1887–1893 (2001).

95. Wald, A. Tests of Statistical Hypotheses Concerning Several Parameters When the Number of Observations is Large. Trans. Am. Math. Soc. 54, 426 (1943).

