## Supplementary material for "Pathogenic tau disrupts the cellular program that maintains neuronal identity": Supplemental Information.docx

**Supplemental Figures**

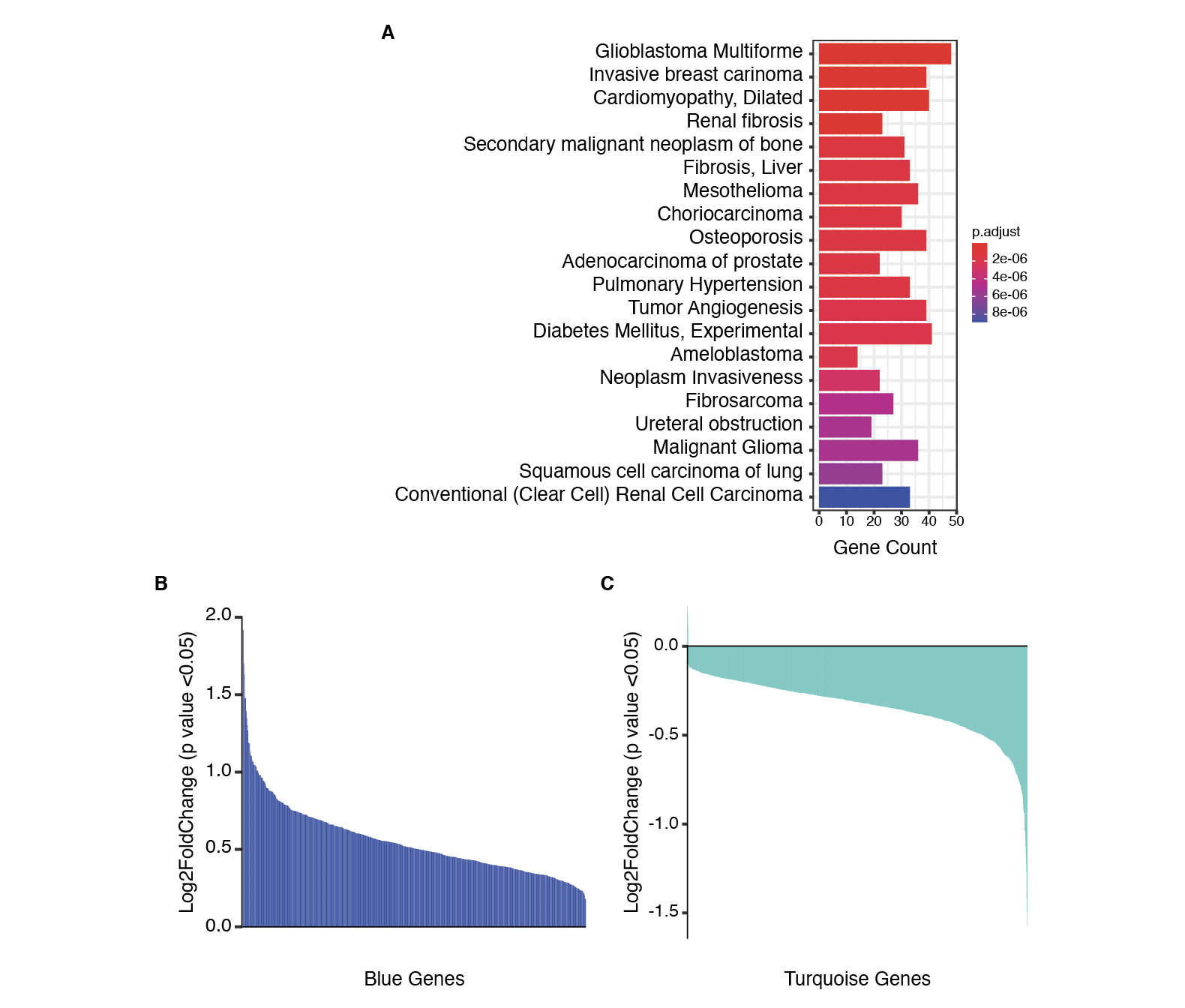

**Figure S1.** **Network analysis of postmortem human Alzheimer’s disease brains identifies a large co-expression module related to EMT and cancer. (A)** Gene-disease association of the blue module in the human network. Bar plot depicts the top 20 most significant DisGeNET terms identified on the y-axis and the number of genes populated in each term on the x-axis. Full table of all DisGeNET terms for each module can be found in Supplementary Table 2. Differentially expressed genes of the blue **(B)** and turquoise **(C)** modules in the human network. Bar plot showing the log2FoldChange of patients with Alzheimer’s disease relative to control for each of the differentially expressed genes from the blue (588 total) and turquoise (1,927 total) modules. Full table of all differentially expressed genes for each module can be found in Supplementary Table 7.

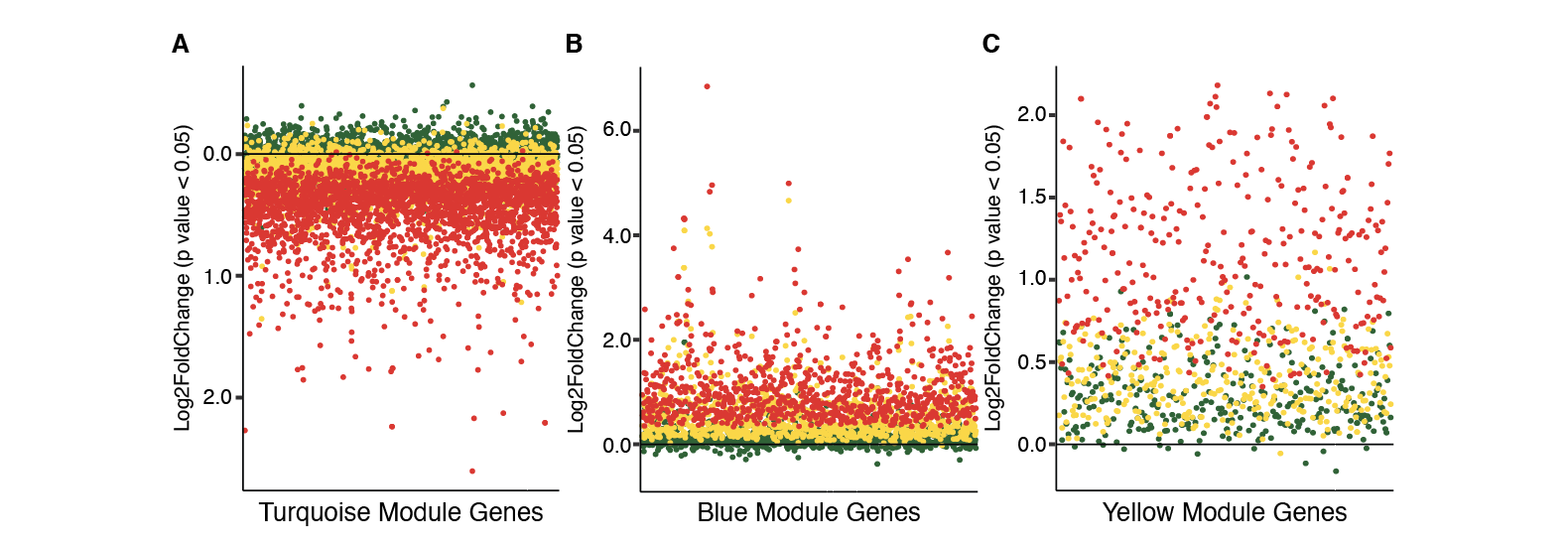

**Figure S2.** **Age-dependent network analysis of tau transgenic mice identifies biological processes in human Alzheimer’s disease that are driven by pathogenic tau and are conserved across disease stage.** Network analysis from female tau transgenic mouse hippocampi show module directionality and magnitude change across aging. Dot plots show all genes within turquoise **(A)**, blue **(B)**, and yellow **(C)** modules and the log2foldChange of tau transgenic mice relative to control associated with each gene at three (green), six (yellow), and nine (red) months. The full table of all modules and the associated genes for each module across aging can be found in Supplementary Table 8.

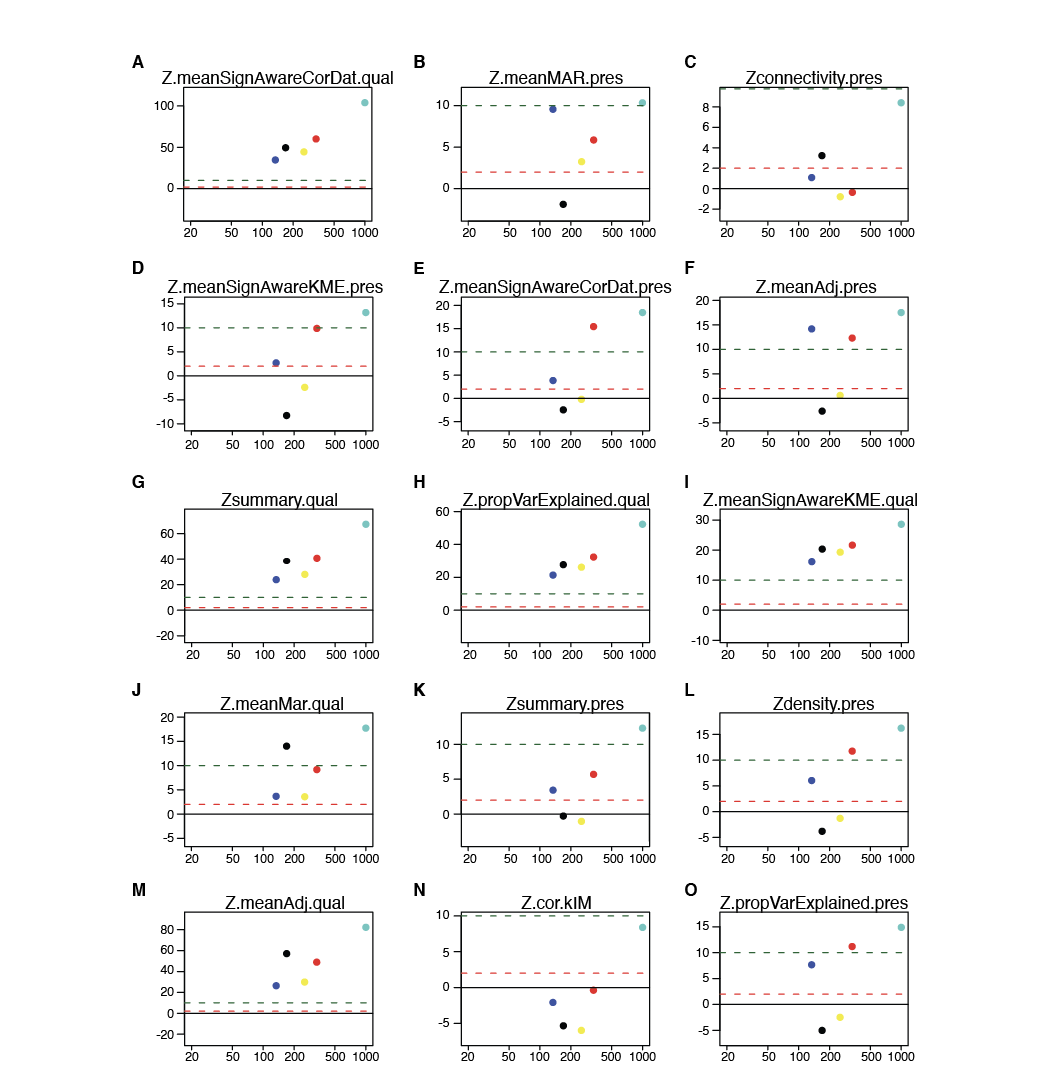

**Figure S3. Different measures of the human network structure are preserved in the mouse network. (A-O)** Preservation statistics calculated for the preservation of the human network (reference) in the mouse (test) network. Each preservation statistic measured a different aspect of the human network preservation and is reported using Z statistics scores. For each graph, module size is labeled along the x-axis and Z scores on the y-axis. Thresholds reflect scoring which dictates that Zscores > 10 are evidence of strong preservation, Zscores > 2 and < 10 are evidence of weak to moderate preservation, and Zscores < 2 are evidence of no module preservation. See Langfelder *et al*. for complete list of definitions and glossary^47^.

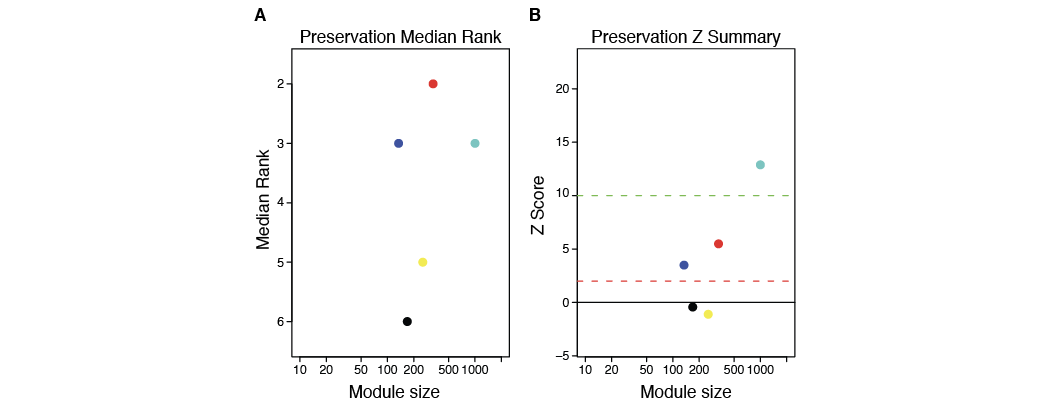

**Figure S4. Turquoise, blue, and red modules of the human network are preserved in the mouse network.** Summary statistics for human module preservation in the tau transgenic mouse network. **(A)** Relative comparison of preservation among multiple modules using *medianRank*. Rankings are indicative of observed preservation statistics such that lower ranking modules tend to be highly preserved. **(B)** Composite of all preservation statistics calculated from module preservation including using summarized statistics from Zdensity and Zconnectivity-based statistics. Thresholds reflect scoring which dictates that Zscores > 10 are evidence of strong preservation, Zscores > 2 and < 10 are evidence of weak to moderate preservation, and Zscores < 2 are no evidence of module preservation. See Langfelder *et al*. for complete list of definitions and glossary^47^.

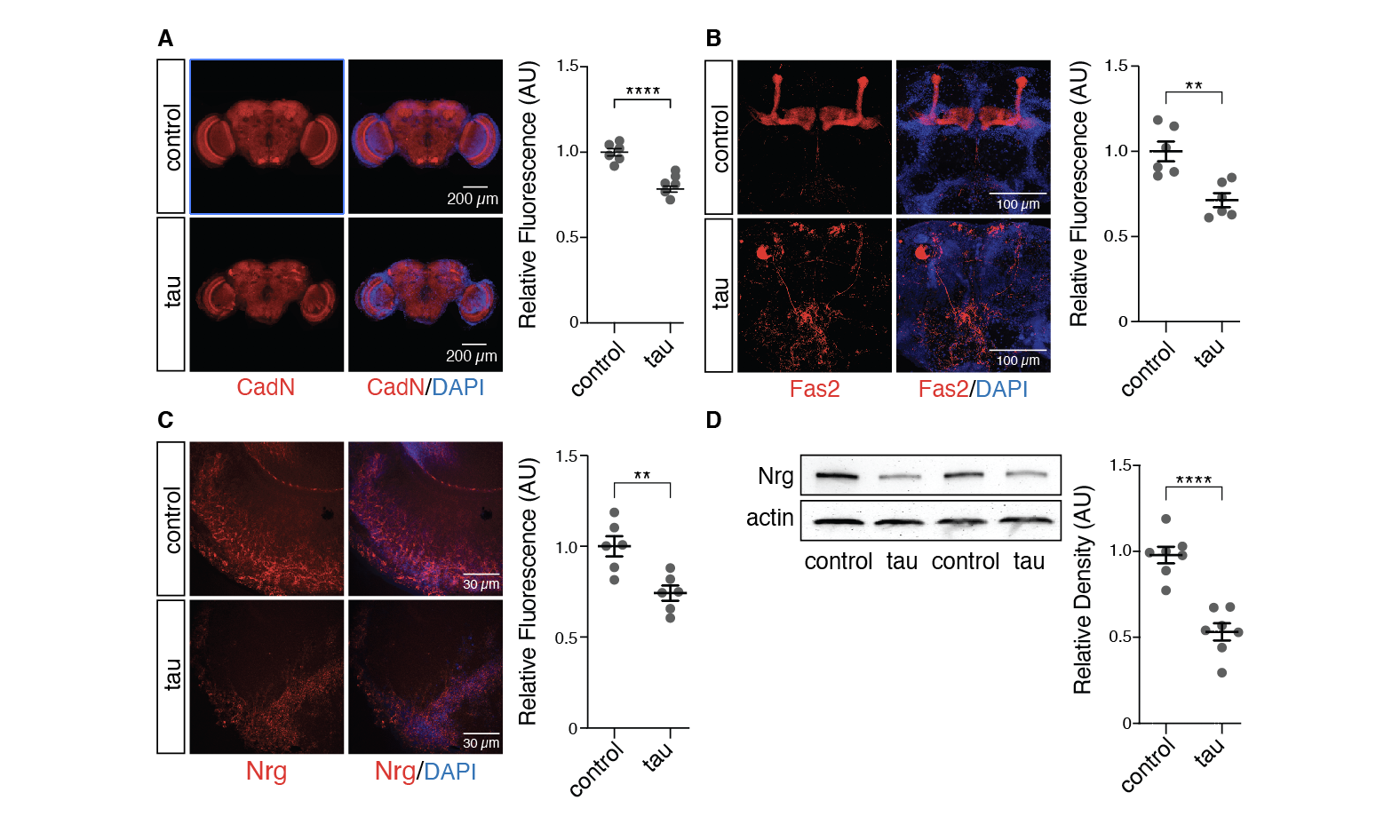

**Figure S5. Brains of tau transgenic *Drosophila* have lower levels of critical adhesion and junction proteins.** Decreased immunofluorescence of **(A)** CadN, **(B)** Fas2, and **(C**) Nrg in brains of tau transgenic *Drosophila*. **(D)** Nrg protein levels are depleted in tau transgenic *Drosophila* based on western blotting. All flies were 10 days old. Values are mean ± s.e.m., n = 6-8 biologically independent replicates per genotype, *P<0.05, **P < 5.0x10^-3^, ****P < 5.0x10^-5^, unpaired, two-tailed Student’s t-test. Full genotypes are listed in Supplementary Table 3.

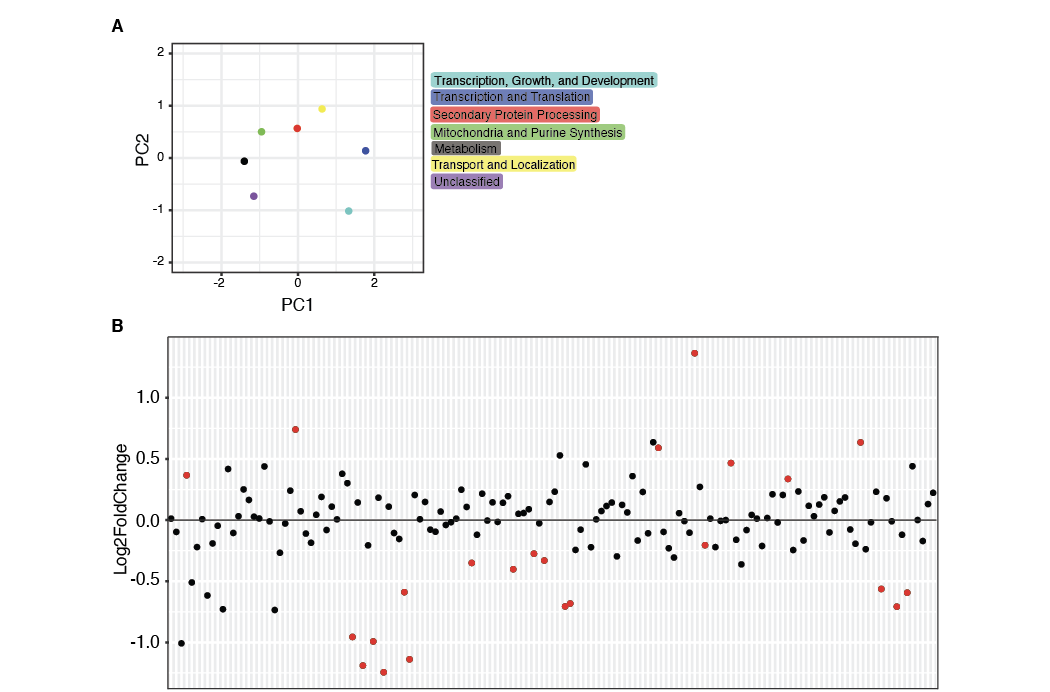

**Figure S6. Pathogenic tau disrupts terminal neuronal differentiation and augments biological processes associated with growth and development. (A)** Multidimensional scaling plot of the first and second principal components for module eigengenes identified by network analysis from *Drosophila* brains. In total, seven modules were identified. The purple (Unclassified) module did not fit the definition (at least 100 genes) used in this study. Colors correspond to module assignment and the adjacent legend provides descriptions based on the most significant biological processes identified from Gene Ontology. A full table of Gene Ontology terms for each module can be found in Supplementary Table 4. **(B)** Dot plot of all prospero target genes identified in *Drosophila* WGCNA. Each dot represents a single gene and depicts log2FoldChange of tau transgenic *Drosophila* relative to control. Red dots signify genes with p values less than 0.05. The full table of gene names and statistics can be found in Supplementary Table 5.

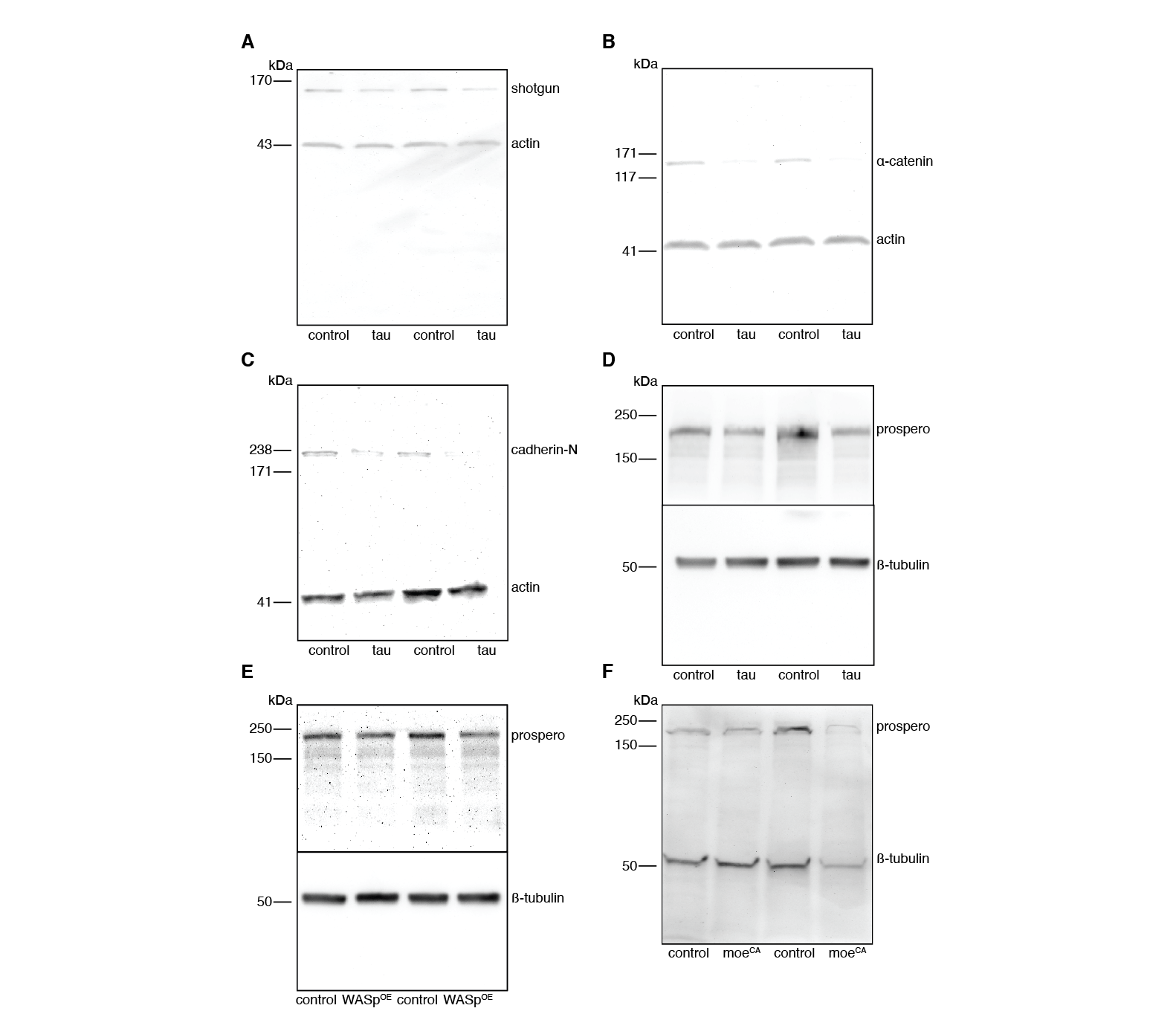

**Figure S7. Full scans of Western blots. (A)** Western blot from Fig. 3B. **(B)** Western blot from Fig 3D. **(C)** Western blot from Fig 3F. **(D)** Western blot from Fig 5B. **(E)** Western blot from Fig 5D. **(F)** Western blot from Fig 5F.
